# lncRNA-ISM1 Promotes Hepatocellular Carcinoma Progression through RBM10-Mediated Alternative Splicing of ISM1 and Akt-S6–Dependent Glucose Metabolic Reprogramming

**DOI:** 10.64898/2026.02.27.708505

**Authors:** Man Li, Deqing Huang, Yuanyuan Ren, Zhen Wang, Yujia Li, Weichen Zuo, Yijie Li, Yanyan Jin, Yuyan Xiong

## Abstract

A low-glucose microenvironment can induce metabolic abnormalities in tumour cells, including hepatocellular carcinoma (HCC) cells, and enhance cancer cell stemness. Isthmin-1 (ISM1) is a recently identified adipokine that promotes glucose uptake and enhances cellular metabolism. While the activity of the ISM1 protein is regulated by glycosylases, its transcriptional and posttranscriptional regulation remain poorly understood. A novel alternatively spliced variant of ISM1 (ISM1-AS) was recently identified. Unlike canonical ISM1, ISM1-AS lacks an AMOP domain, a key structural element required for ISM1 function, suggesting the loss of its metabolic regulatory activity. In this study, we found that ISM1 expression was significantly reduced in HCC tissues and correlated with poor prognosis. Functional assays revealed that ISM1 overexpression markedly suppressed HCC cell proliferation and invasion, whereas ISM1-AS overexpression had the opposite effect. Importantly, ISM1 co-overexpression attenuated the oncogenic effects of ISM1-AS. Knockdown of the antisense transcript lncRNA-ISM1 reduced ISM1-AS expression while increasing ISM1 expression, thereby suppressing HCC proliferation and migration. Mechanistically, lncRNA-ISM1 regulated ISM1 alternative splicing by interacting with RBM10, thereby altering the balance between ISM1-AS and ISM1. This shift activated the Akt-S6 signalling pathway, promoting glycolysis and HCC progression. In vivo experiments further confirmed that the lncRNA-ISM1/ISM1-AS/ISM1 axis drives tumour growth via Akt-S6 activation. Our findings demonstrate that lncRNA-ISM1 promotes HCC progression through the RBM10-mediated alternative splicing of ISM1 and activation of the Akt-S6 signalling pathway, highlighting its potential as a therapeutic target for HCC.

## 1. Introduction

HCC has become the sixth most common cancer globally and the fourth most common cancer in China, accounting for up to 75–85% of primary liver cancers [1–3]. Currently, surgery remains the preferred treatment for HCC patients. Unfortunately, the mortality rate has significantly decreased in only some carefully treated HCC patients. In other patients, the rates of tumour metastasis and recurrence are still very high, and even after surgery, the lifespan of these patients is relatively short [4]. Despite significant progress in clinical and experimental cancer treatments, owing to the high rates of postoperative tumour recurrence and metastasis, the overall prognosis of HCC patients is poor [5, 6]. Previous studies have shown that the progression of HCC may involve multiple steps involving the participation of multiple factors and altered expression of multiple genes, but the specific molecular mechanisms of HCC metastasis remain unclear [7]. Therefore, a better understanding of the underlying molecular mechanisms of HCC metastasis is highly important for the targeted treatment and prognostic evaluation of HCC.

The metabolic patterns of tumours are significantly different from those of normal cells, and glucose metabolism is an important pathway for tumour cells to obtain energy and biosynthetic precursors [8, 9]. In recent years, studies have shown that many cancer cells exhibit significant reprogramming in glucose metabolism. In particular, in a hypoxic environment, tumour cells tend to increase glycolysis to meet their needs for rapid proliferation; this phenomenon is known as the “Warburg effect” [10, 11]. Through this metabolic adaptation, tumour cells can rapidly generate ATP and accumulate key metabolites such as lactate and α-ketoglutarate. These metabolites not only provide energy for cells but also serve as biosynthetic precursors for the proliferation, invasion, and metastasis of tumour cells [12]. Aerobic glycolysis was first discovered in HCC and is involved in the regulation of proliferation, immune escape, invasion, metastasis, angiogenesis, and drug resistance in liver cancer [1]. Glycolytic pathway activation in hepatocellular carcinoma cells is related to various factors, including the increased expression of hypoxia-inducible factors (HIFs), the upregulation of glucose transporter (GLUT) expression, and the activation of rate-limiting enzymes such as HK2, PFK1, and PKM2. In this context, liver cancer is among the cancers with high incidence and mortality rates globally [13]. The liver plays a key role in whole-body metabolism, and hepatocellular carcinoma cells often promote their own growth and adapt to changes in the microenvironment by altering the glucose metabolism pathway. Therefore, the study of glucose metabolism in the liver is particularly important [14]. Multiple studies have shown that the expression of key enzymes related to glucose metabolism in hepatocellular carcinoma cells is abnormal and that the activation of signalling pathways such as the PI3K/Akt/mTOR pathway further promotes glycolysis [15]. Metabolic changes in hepatocellular carcinoma cells increase their survival advantages and accelerate disease progression. Studying these glucose metabolism mechanisms not only helps to elucidate the biological basis of liver cancer but also provides key targets for the development of new therapies and early diagnosis.

Isthmin-1 (ISM1) is a novel adipokine that was initially discovered in the embryos of *Xenopus laevis* and plays a role in embryonic and early brain development [16, 17]. In recent years, increasing evidence has shown that the abnormal expression of ISM1 can affect the biological behaviour of cancer [18]. For example, the downregulation of ISM1 expression promotes the proliferation of colon adenocarcinoma cells, inhibits apoptosis, and promotes the expression of cyclins [19]. ISM1 deficiency leads to enhanced glycolysis and increased lactate production in hepatocellular carcinoma cells, thus promoting tumour progression [20]. In gastric cancer, lncRNA H19 can promote the occurrence, development, and metastasis of gastric cancer by regulating the expression level of ISM1 [21]. In addition, has-circ-0091570 inhibits the progression of hepatocellular carcinoma cells by acting as a ceRNA to competitively bind to miR-1307 and regulate the expression of ISM1 [22]. However, to a large extent, whether ISM1 in HCC is regulated by other factors remains unclear.

Recent studies have shown that alternative splicing plays an important role in cancer and interacts with other molecular mechanisms to promote both the development and progression of cancer [23]. Alternative splicing generates multiple mRNA splicing isoforms from precursor mRNA (pre-mRNA) through the selective removal of introns and retention of exons [24]. Transcripts of almost all human protein-coding genes undergo one or more forms of alternative splicing [25], which increases the diversity of gene phenotypes and protein functions in eukaryotes [26, 27]. Alternative splicing is among the important mechanisms of gene expression regulation. In hepatocellular carcinoma, determining whether the expression and function of ISM1 are affected by alternative splicing (ISM1-AS) is important.

Noncoding RNAs (ncRNAs) are a class of abundant RNAs that do not encode proteins and play important roles in cancer by regulating alternative splicing, affecting various cellular signalling pathways, and promoting or inhibiting cancer progression [28]. In recent years, the functions of long noncoding RNAs (lncRNAs) in HCC have received much attention. They participate in transcriptional and posttranscriptional regulation through various mechanisms (such as molecular scaffolding, RNA–RNA duplex formation, and the regulation of protein recruitment) [29–32]. Although the roles of various lncRNAs in HCC have been widely studied [33], whether the ISM1 antisense transcript (lncRNA-ISM1) regulates the alternative splicing of ISM1 mRNA and affects HCC cell glycolysis and HCC progression remains unknown.

In this study, we revealed that lncRNA-ISM1 affects the expression of ISM1 by transcriptionally regulating ISM1-AS, which in turn activates the Akt-S6 signalling pathway and promotes glycolysis and cell progression in HCC cells. These findings elucidate the mechanism of action of the lncRNA-ISM1/ISM1-AS/ISM1 axis in tumour progression and suggest that lncRNA-ISM1 may serve as a potential target for the diagnosis and treatment of HCC.

## 2. Materials and Methods

### 2.1 Reagents

The following reagents were purchased or obtained from the following sources: rabbit anti-ISM1 antibody (PA5-24968, Invitrogen, USA), rabbit anti-ISM1-AS antibody (MR-YT-20221208-001, Mabioway, China), rabbit anti-GAPDH antibody (#2118s, CST, USA), rabbit anti-total AKT (#4685s, CST, USA), rabbit anti-phospho-AKT (Ser 473) (#4060t, CST, USA), rabbit anti-total S6 (#2217s, CST, USA), rabbit phospho-S6 (Ser 235/236) (#4858t, CST, USA), mouse anti-GLUT4 antibody (66846-1-Ig, Proteintech, China), rabbit anti-HK2 antibody (22029-1-AP, Proteintech, China), rabbit anti-LDH antibody (14546-1-AP, Proteintech, China), rabbit anti-PKM2 antibody (15822-1-AP, Proteintech, China), and rabbit anti-RBM10 (84104-2-RR, Proteintech, China). All cell culture media and materials were purchased from Gibco (Waltham, MA).

### 2.2 Tissue samples

The tissues used in this study, including liver cancer tissues and adjacent normal tissues, were collected during surgical resection at Xi’an Third Hospital, Affiliated Hospital of Northwest University. All patients who participated in this study signed informed consent forms, and the use of tissue samples in this study was reviewed and approved by the Ethics Committee of Xi’an Third Hospital, Affiliated Hospital of Northwest University.

### 2.3 Cell culture

The HCC cell lines MHCC-97H and Hep3B used in this study were purchased from the Cell Bank of the Chinese Academy of Sciences (Shanghai, China). All cells were cultured in a 37°C incubator with 5% CO_2_ in Dulbecco’s modified Eagle’s medium (Sigma, USA) supplemented with 10% foetal bovine serum (Sigma, USA) and 100 μg/mL penicillin‒streptomycin. All the cell lines were tested for mycoplasma contamination. After being confirmed to be in good condition through cell viability assays (such as trypan blue staining) and microscopic morphological observations, the cells were used in subsequent experiments.

### 2.4 Plasmid construction and cell transfection

The ISM1 overexpression vector plasmid (PCMV-ISM1), ISM1-AS overexpression vector plasmid (PCMV-ISM1-AS), lncRNA-ISM1 overexpression vector plasmid (PCMV-LncRNA-ISM1) and empty vector plasmid were purchased from GenePharma (Shanghai, China). The ISM1 siRNA, ISM1-AS siRNA, lncRNA-ISM1 siRNA, RBM10 siRNA and negative control sequence were designed and synthesized by GenePharma and cloned into the LV3 (H1/GFP&Puro) vector. The vectors were transfected into HCC cell lines using Lipofectamine 2000 in accordance with the manufacturer’s instructions. The cells were screened with puromycin for 14 days to obtain stably transfected cell lines.

### 2.5 Quantitative real-time PCR

Total RNA was extracted from the cell lines using TRIzol reagent (Thermo Fisher, Waltham, MA, USA), and the total RNA was reverse transcribed into cDNA using a PrimeScript^TM^ kit (TaKaRa, Dalian, China). The cDNA was amplified by SYBR green-based qRT‒PCR on a Bio-Rad CFX96 RealTime PCR system (Bio-Rad, US). The procedure was as follows: predenaturation at 95°C for 30 s, followed by 40 cycles of 95°C for 5 s, 55°C for 30 s, and 72°C for 30 s. GAPDH was used as an internal reference. The relative gene expression levels were calculated using the 2^−△△Ct^ method. The names and sequences of the primers used in this study are as follows (5’-3’):

ISM1-F: ATTATCTGCAAGGGTGACTGG,
ISM1-R: CCTCTTGGAACTGCTTGATGTA;
ISM1-AS-F: GACAAATGGGAGTGGCTGTA,
ISM1-AS-R: AAGTGTTGGTGAGGATGTGTAG;
LncRNA-ISM1-F: TGGAGAACAATCTGTGGGATAG,
LncRNA-ISM1-R: CAGGGTACATGGTGTAGAAGAG;
TUG1-F: GCAAACTGAGGATGCTCCATCC,
TUG1-R: TACCAGGTCTGTAGGCTGATGG;
BMP2-F: TGTATCGCAGGCACTCAGGTCA,
BMP2-R: CCACTCGTTTCTGGTAGTTCTTC;
ATM-F: TGTTCCAGGACACGAAGGGAGA,
ATM-R: CAGGGTTCTCAGCACTATGGGA;
RBM10-F: GGCATCTACCAACAATCAGCCG,
RBM10-R: GGAGAGCAGAACTAGGATGGGT;
GAPDH-F: TCTCCAACTCTCAGCCCACCAA,
GAPDH-R: CCTGCGATGCTGGACTTGACCT.

### 2.6 Western blotting assay

Cells or tissue samples were placed in RIPA lysis buffer containing protein phosphatase inhibitors and phenylmethylsulfonyl fluoride (PMSF) and lysed on ice for 30 minutes to extract proteins. Then, the proteins were separated by SDS‒polyacrylamide gel electrophoresis and transferred onto a 0.2-μm PVDF membrane (Millipore, Bedford, MA). After blocking with 5% skim milk at room temperature for 2 hours, the membrane was incubated with a primary antibody overnight at 4°C. Then, the membrane was incubated with the corresponding secondary antibody at room temperature for 1 hour and washed three times in TBST. The protein bands were visualized using a chemiluminescence imaging system (GE Healthcare, Chicago, USA).

### 2.7 EdU proliferation assay

Cell proliferation was determined using a BeyoClick™ EdU-488 Cell Proliferation Assay Kit (C0071S; Beyotime, China). The cells were seeded at a density of 3×10⁴ cells per well in a 24-well plate, with five replicate wells for each group. After the cells were cultured for 24 hours, 250 μl of EdU working solution was added to each well, and the plate was incubated in the incubator for 2 hours. After incubation, the culture medium was removed, and the cells were fixed with 4% paraformaldehyde at room temperature for 15 minutes. Then, 100 μl of the click reaction solution was added to each well, and the plate was incubated in the dark at room temperature for 30 minutes. Finally, Hoechst stock solution was diluted with PBS at a ratio of 1:1000, and 100 μl of the diluted solution was added to each well. The cells were incubated in the dark at room temperature for 10 minutes for nuclear staining, and cell proliferation status was assessed under a fluorescence microscope.

### 2.8 Colony formation assay

Cells were seeded at a density of 500 cells per well in a 6-well plate and cultured in medium supplemented with 10% FBS for approximately 2 weeks until colony formation was observed under a microscope. Then, the cells were fixed in 4% paraformaldehyde for 30–60 minutes, stained with a 0.2% crystal violet solution (Sigma, St. Louis, MO, USA) for 15 minutes, and finally counted using ImageJ.

### 2.9 Transwell assay

Tumour cell migration was assessed using Transwell chambers with an 8.0-μm pore size (Becton Dickinson, Bedford, MA, USA). In the upper chamber, 200 μl of serum-free medium containing 5×10⁵ cells was added, and 600 μl of medium containing 15% FBS was added to the lower chamber. After the cells were cultured for 24 hours, the noninvading cells in the upper chamber were removed with a cotton swab. The invading cells on the reverse side were fixed with 4% methanol and then stained with a 0.25% crystal violet solution. Five random fields of view were selected from each chamber for imaging and counting, and a statistical graph was drawn.

### 2.10 calorimetry system

The contents of glucose, lactate, and ATP in cells were determined using a glucose content assay kit, lactate dehydrogenase assay kit, and ATP content assay kit (mlbio, Shanghai, China), respectively. The cells were collected in a centrifuge tube, and after centrifugation, the supernatant was discarded. The appropriate amount of the corresponding extraction solution was added, and the cells were disrupted by sonication. After centrifugation, the supernatant was removed, and the assays were carried out in strict accordance with the instructions of the kits.

### 2.11 RNA immunoprecipitation

Cells were washed with precooled PBS, and lysis buffer containing a protease inhibitor was added. The cells were lysed on ice for 30 minutes. The composition of the lysis buffer was 40 mM HEPES (pH 7.5), 120 mM NaCl, 1 mM EDTA, 10 mM pyrophosphate, 10 mM glycerophosphate, and 0.3% CHAPS. After the cells were scraped, they were centrifuged at 10,000 ×g for 15 minutes at 4°C to remove cell debris. The supernatant was collected and transferred to a new 1.5 mL centrifuge tube. One percent of the supernatant sample was mixed with 490 μl of TRIzol as an RNA input control. The lysate was precleared using protein A/G magnetic beads and incubated with a primary antibody at 4°C for 2 hours. Then, 20 μl of protein A/G magnetic beads were added, and the mixture was gently rotated and incubated overnight at 4°C. The immunoprecipitates were collected by centrifugation at 2500 rpm for 5 minutes at 4°C and then were washed 3–4 times with wash buffer (containing 50 mM HEPES (pH 7.5), 40 mM NaCl, and 2 mM EDTA). Finally, the magnetic beads containing the immunoprecipitated samples were resuspended in 500 μl of TRIzol for qRT‒PCR analysis.

### 2.12 Animal studies

Six-week-old male BALB/c nude mice were purchased from Beijing Vital River Laboratory Animal Technology Co., Ltd. (Beijing, China) and were raised in a constant-temperature and -humidity environment. Animal experiments were conducted in accordance with the guidelines of the Animal Care and Use Committee of Northwest University (Xi’an, Shaanxi, China; Approval No.: NWU-AWC-20230102M; January 18, 2023) and followed the guidelines of the National Institutes of Health (NIH) in the United States. In the tumorigenesis experiments, 36 nude mice were randomly divided into 6 groups. MHCC-97H cells stably expressing the lentiviral vector or lentiviral interference vector (2×10⁶ cells per mouse, 0.1 mL of PBS) were subcutaneously inoculated into the flanks of 6-week-old male BALB/c nude mice (n = 6 per group). The status of the mice was monitored daily, and the volume of the subcutaneous tumours was measured using a Vernier calliper twice a week. The formula for calculating tumour volume (mm³) was as follows: Volume = (Length × Width²)/2. Four weeks later, all the mice were euthanized. The tumour tissues were collected, and their weights were recorded. The tissues were subsequently stored at −80°C for subsequent analysis.

### 2.13 Immunohistochemistry (IHC)

Tumour tissues were fixed in 4% paraformaldehyde, embedded in paraffin, and then cut into sections (4 μm thick). The sections were dewaxed with xylene and rehydrated through a gradient of ethanol solutions. The sections were incubated with a 3% H₂O₂ solution at room temperature for 10 minutes to quench endogenous peroxidase activity, after which they were washed three times with PBS. Nonspecific antigens were blocked with PBS containing 5% BSA for 30 minutes. The sections were boiled in 10 mM citric acid buffer for 30 minutes for antigen retrieval. After the samples were allowed to cool to room temperature, the corresponding primary antibody was added, and the sections were incubated overnight at 4°C and then incubated with an HRP-labelled secondary antibody. After dehydration and clearing, the sections were stained with DAB substrate, counterstained with haematoxylin, and sealed with neutral gum. Finally, the sections were observed and photographed through a DM3000 microscope (Leica, Wetzlar, Germany). The images were analysed using ImageJ software.

### 2.14 Immunofluorescence staining

Tissue sections were washed three times with PBS (5 minutes each) to remove OCT embedding medium. The sections were subsequently permeabilized in 0.2% Triton X-100 for 20 minutes and blocked with PBS containing 5% BSA for 1 hour. After the blocking solution was removed, a primary antibody was added, and the sections were incubated overnight at 4°C. Then, the sections were incubated with a secondary antibody in the dark at room temperature for 1 hour. The cells were incubated with DAPI in the dark for 20 minutes to label the cell nuclei. Finally, the sections were sealed with anti-fluorescence quenching mounting medium, and images were collected using a fluorescence microscope. All the images were randomly obtained from different fields of view, and the relative fluorescence intensity was analysed using ImageJ software.

### 2.15 Statistical analysis

Statistical analysis was performed using GraphPad Prism 8.0 software. All the quantitative experiments were independently repeated three times. The data are expressed as the mean ± standard deviation. A t test was used for comparisons between two groups, and one-way analysis of variance (ANOVA) was used for comparisons among multiple groups to determine statistical significance. A P value less than 0.05 was considered to indicate statistical significance, and the significance is indicated as follows: *P < 0.05, **P < 0.01, ***P < 0.001, and ****P < 0.0001; n.s. indicates no significant difference.

## 3. Results

### 3.1 ISM1 expression is aberrantly downregulated in HCC tissues and cells

To investigate the expression of ISM1 in patients with hepatocellular carcinoma (HCC), we first obtained gene expression data for ISM1 from the TCGA database. The results revealed that the expression of ISM1 was significantly lower in HCC tumour tissues than in normal tissues **(Fig. 1A)**. To evaluate the prognostic value of ISM1 in liver cancer, Kaplan–Meier (K–M) survival curve analysis indicated that lower levels of ISM1 expression were associated with a reduced overall survival rate **(Fig. 1B)**. Immunohistochemical analysis further confirmed that the protein expression level of ISM1 was significantly lower in HCC tissues than in adjacent noncancer tissues **(Fig. 1C–D)**. In addition, compared with those in the human normal liver cell line LO2, the mRNA **(Fig. 1E)** and protein **(Fig. 1F)** expression levels of ISM1 were abnormally downregulated in multiple HCC cell lines (including Hep3B, Huh-7, MHCC-97H, and HepG2). Since the downregulation of ISM1 expression was more significant in MHCC-97H and Hep3B cells, these two cell lines were selected for subsequent in vitro experiments. These data suggest that ISM1 may inhibit the progression of HCC.

**Fig. 1.**
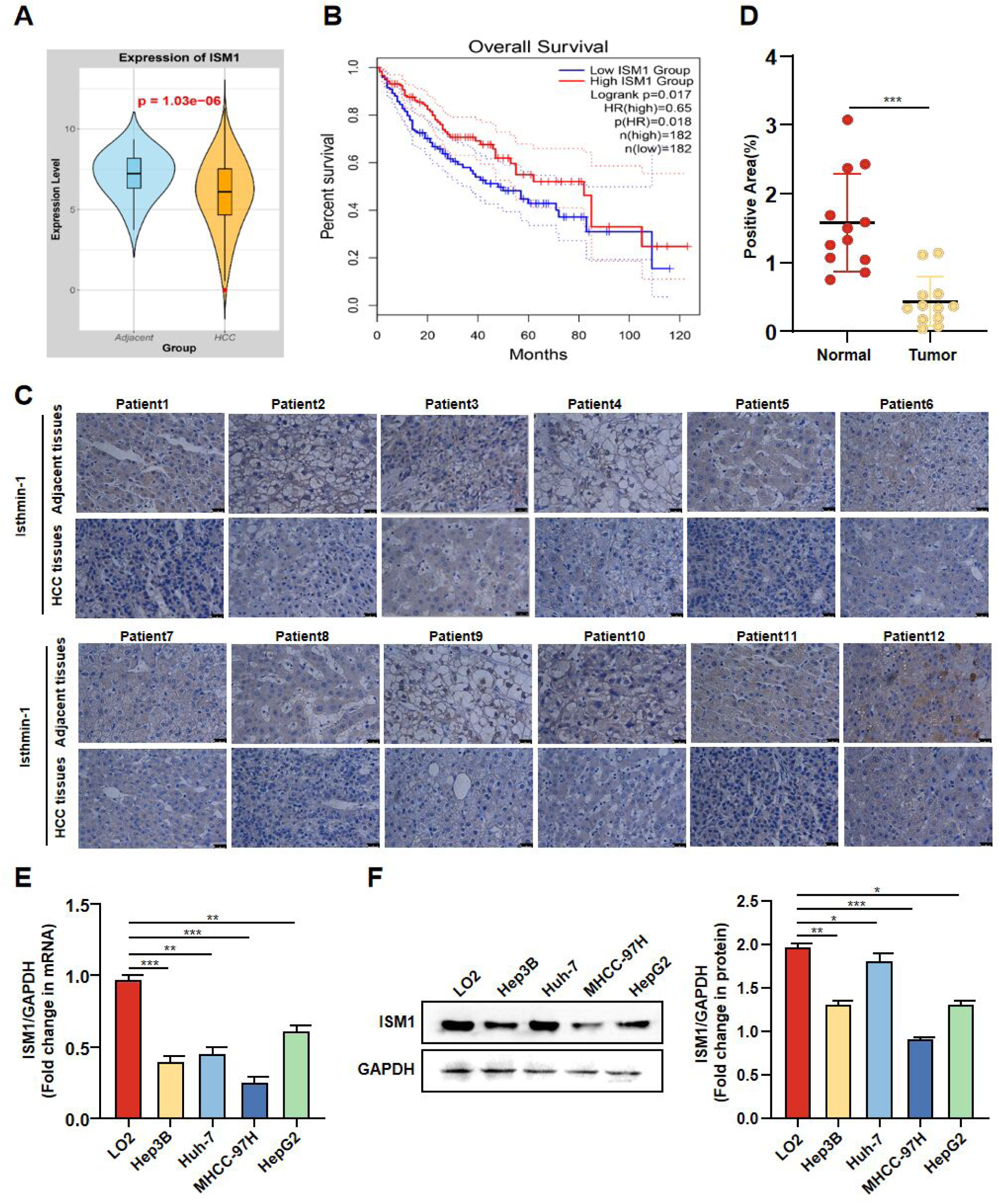
ISM1 expression is aberrantly downregulated in HCC tissues and cells. **(A)** The relative expression levels of ISM1 in HCC and paired normal samples from the TCGA database. **(B)** A Kaplan‒Meier (K‒M) curve from the TCGA database was used to evaluate the prognostic value of ISM1 in HCC. (C–D) ISM1 expressionin liver cancer tissues and the corresponding adjacent noncancer tissues was assessed by immunohistochemistry (IHC). **(E)** The mRNA levels of ISM1 in LO2, Hep3B, Huh-7, MHCC-97H, and HepG2 cells were assessed by qPCR. **(F)** ISM1 protein levels were assessed by Western blot. Statistical differences were identified by unpaired Student’s t test. The data are presented as the mean ± SEM. n=3-12. * *p* < 0.05, ** *p* < 0.01, and *** *p* < 0.001. Scale bar = 100 μm.

### 3.2 ISM1 inhibits HCC cell proliferation and invasion

Although ISM1 expression is significantly downregulated in HCC cells, whether this downregulation affects the proliferation and invasion of HCC cells remains unclear. To explore the biological function of ISM1 in HCC progression, we transfected MHCC-97H and Hep3B cells with ISM1-overexpressing lentivirus (0.5 μg/mL and 1 μg/mL). Then, the cells were selected with puromycin for 14 days to obtain cell lines stably transfected with ISM1. The transfection efficiency was verified by Western blotting **(Fig. 2A-B)**. EdU-based proliferation and colony formation assays revealed that ISM1 overexpression significantly inhibited the proliferation and colony formation abilities of MHCC-97H and Hep3B cells **(Fig. 2C–F)**. To investigate the role of ISM1 in HCC invasion, we conducted a Transwell invasion assay. As shown in **Fig. 2G–H**, ISM1 overexpression significantly reduced the invasion ability of MHCC-97H and Hep3B cells. In addition, we effectively knocked down the expression of ISM1 in MHCC-97H and Hep3B cells using siRNA (30 nM and 60 nM) **(Fig. S1A–B)**, and subsequent experiments investigated the tumour suppressor role of ISM1 in regulating the proliferation and invasion of HCC cells. ISM1 knockdown significantly enhanced the proliferation **(Fig. S1C-D)** and colony formation ability **(Fig. S1E-F)** of MHCC-97H and Hep3B cells. Moreover, after ISM1 knockdown, the invasive ability of the cells significantly increased **(Fig. S1G-H)**. These results indicate that ISM1 plays a tumour suppressor role in inhibiting the proliferation and invasion of HCC cells.

**Fig. 2.**
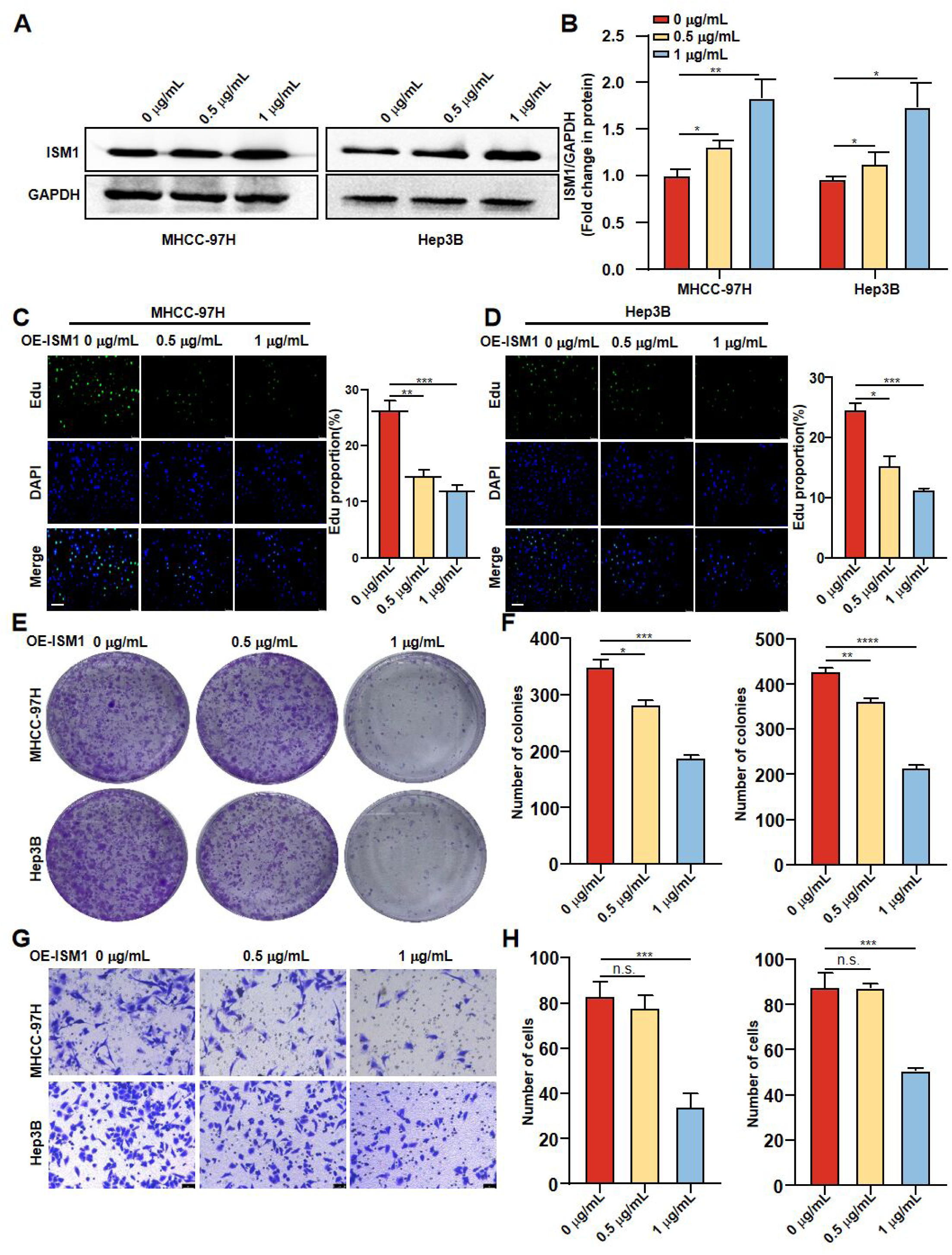
ISM1 overexpression inhibits the proliferation and invasion of HCC cells. MHCC-97H cells and Hep3B cells were infected with an ISM1-overexpressing lentivirus (0.5 μg/mL and 1 μg/mL). **(A-B)** The protein level of ISM1 was assessed by Western blotting. **(C–D)** EdU-based proliferation and **(E–F)** colony formation assays were used to assess cell proliferation ability. **(G–H)** Transwell assays were used to evaluate the invasion ability of MHCC-97H cells and Hep3B cells. Statistical differences were identified by unpaired t test. The data are presented as the mean ± standard error, n = 3. * *p* < 0.05, ** *p* < 0.01, *** *p* < 0.001, and **** *p* < 0.0001; n.s., not significant. Scale bar = 100 μm.

### 3.3 ISM1 promotes the proliferation and invasion of HCC cells

A large body of evidence suggests that alternative splicing is among the important mechanisms for gene expression regulation[27, 34]. In 2022, a new alternative splice variant of ISM1 (ISM1 - AS) was discovered in GenBank. Unlike the classic ISM1 protein, the protein encoded by this alternative splice variant lacks an AMOP domain, which is an important structural basis for the function of ISM1. This prompted us to explore whether ISM1-AS is involved in the tumour-suppressive effect of ISM1 on HCC. Interestingly, after ISM1-AS was overexpressed in MHCC-97H and Hep3B cells **(Fig. S2A-B)**, the proliferation ability of HCC cells increased; these results were verified by EdU proliferation and colony formation assays **(Fig. S2C-F)**. Moreover, the results of Transwell invasion assays also revealed that the overexpression of ISM1-AS significantly enhanced the invasion ability of HCC cells **(Fig. S2G-H)**. Next, we overexpressed ISM1 via lentivirus in MHCC-97H and Hep3B cells overexpressing ISM1-AS **(Fig. 3A–B)**. Strikingly, we found that the overexpression of ISM1 ameliorated the promoting effect of ISM1-AS overexpression on tumour cell proliferation, and these results were verified in MHCC-97H and Hep3B cells through EdU proliferation and colony formation assays (**Fig.** 3C–F). Moreover, Transwell invasion assays revealed that ISM1 expression significantly increased the ability of ISM1-AS to promote the invasion of HCC cells **(Fig. 3G–H)**. These results provide evidence that ISM1 promotes the proliferation and invasion of HCC cells through ISM1-AS overexpression.

**Fig. 3.**
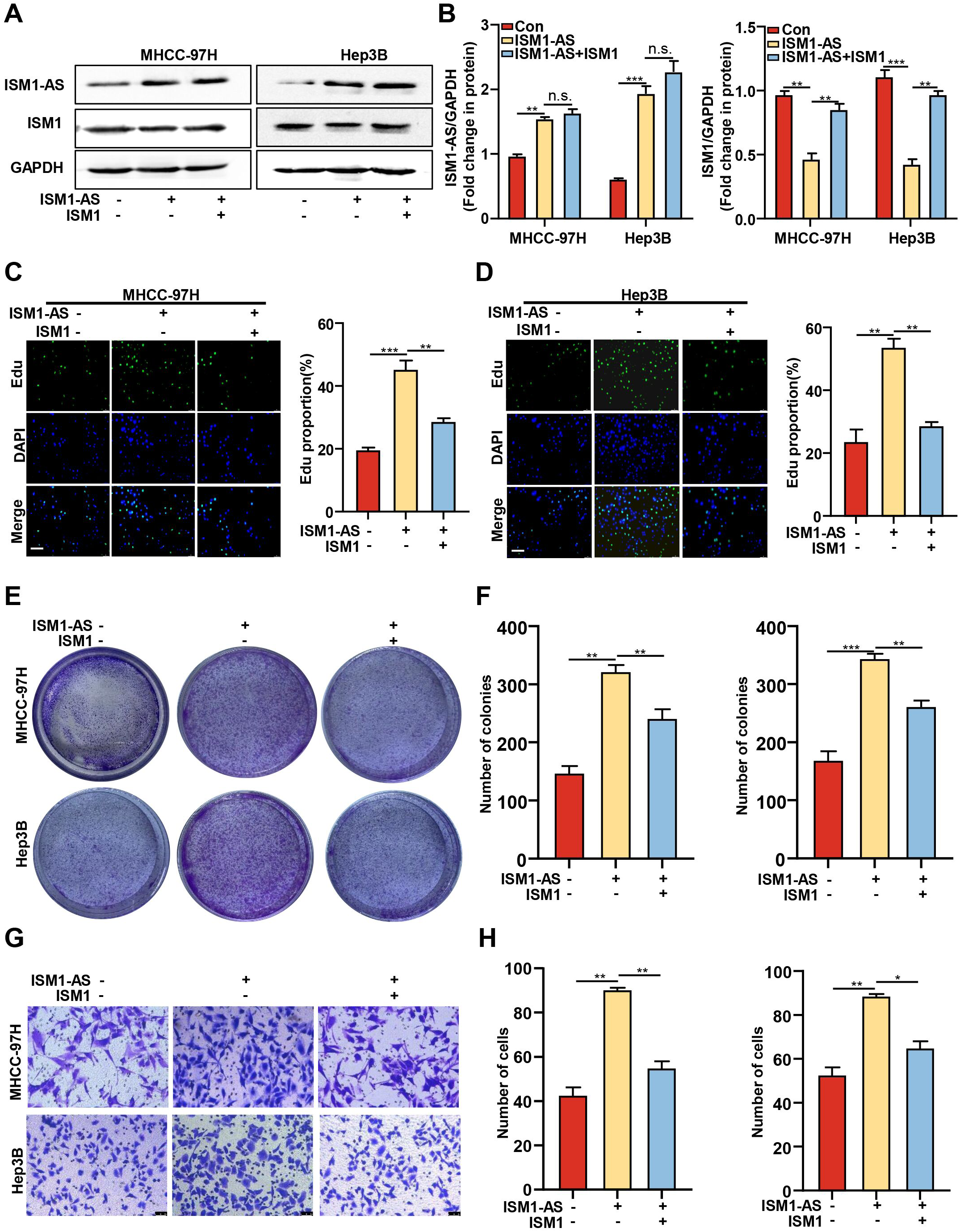
ISM1 ameliorates the promoting effect of ISM1-AS overexpression on the proliferation and invasion of HCC cells. ISM1-overexpressing lentivirus was transfected into MHCC-97H and Hep3B cells that had been transfected with an ISM1-AS-overexpressing lentivirus, and transfection was conducted for 24 hours. **(A–B)** The protein levels of ISM1-AS and ISM1 in the cells were assessed by Western blotting. **(C–D)** EdU-based proliferation and **(E–F)** colony formation assays were used to evaluate the proliferation ability of the cells. **(G–H)** Transwell migration assays were performed to assess the invasive ability of MHCC-97H and Hep3B cells. Unpaired Student’s t test was used to detect significant differences. The data are presented as the mean ± standard error of the mean (mean ± SEM). n = 3. * *p* < 0.05, ** *p* < 0.01, *** *p* < 0.001; n.s., not significant. Scale bar = 100 μm.

### 3.4 lncRNA-ISM1 regulates the proliferation and invasion of HCC cells through the ISM1-AS/ISM1 axis

An antisense transcript refers to an RNA molecule transcribed from the antisense strand of a gene, and such transcripts have been shown to affect the alternative splicing of mRNA and tumour progression [35]. To investigate the effect of the ISM1 antisense transcript (lncRNA-ISM1) on ISM1-AS and ISM1, we used three pairs of siRNAs to knockdown the expression of lncRNA-ISM1, and knockdown efficiency was verified by qPCR **(Fig. 4A)**. Then, the expression of lncRNA-ISM1 was knocked down by siRNA in MHCC-97H and Hep3B cells, and knockdown efficiency was assessed by qPCR and Western blot analysis. The knockdown of lncRNA-ISM1 led to a decrease in ISM1-AS and an increase in ISM1 (**Fig.** 4B–D). Moreover, EdU proliferation and colony formation assays revealed that the knockdown of lncRNA-ISM1 inhibited the proliferation of HCC cells **(Fig. 4E–G)**. Transwell invasion assays also demonstrated that the knockdown of lncRNA-ISM1 significantly inhibited the invasion of HCC cells **(Fig. 4H)**. In contrast, the overexpression of lncRNA-ISM1 in MHCC-97H and Hep3B cells **(Fig. S3A-D)** led to an increase in ISM1-AS and a decrease in ISM1 and promoted the proliferation of HCC cells, as verified by EdU proliferation and colony formation assays **(Fig. S3E-G)**. Moreover, Transwell invasion assays revealed that the overexpression of lncRNA-ISM1 promoted the invasion of HCC cells **(Fig. S3H)**. These data provide strong evidence that lncRNA-ISM1 exerts its pro-carcinogenic effect by regulating the ISM1-AS/ISM1 axis.

**Fig. 4.**
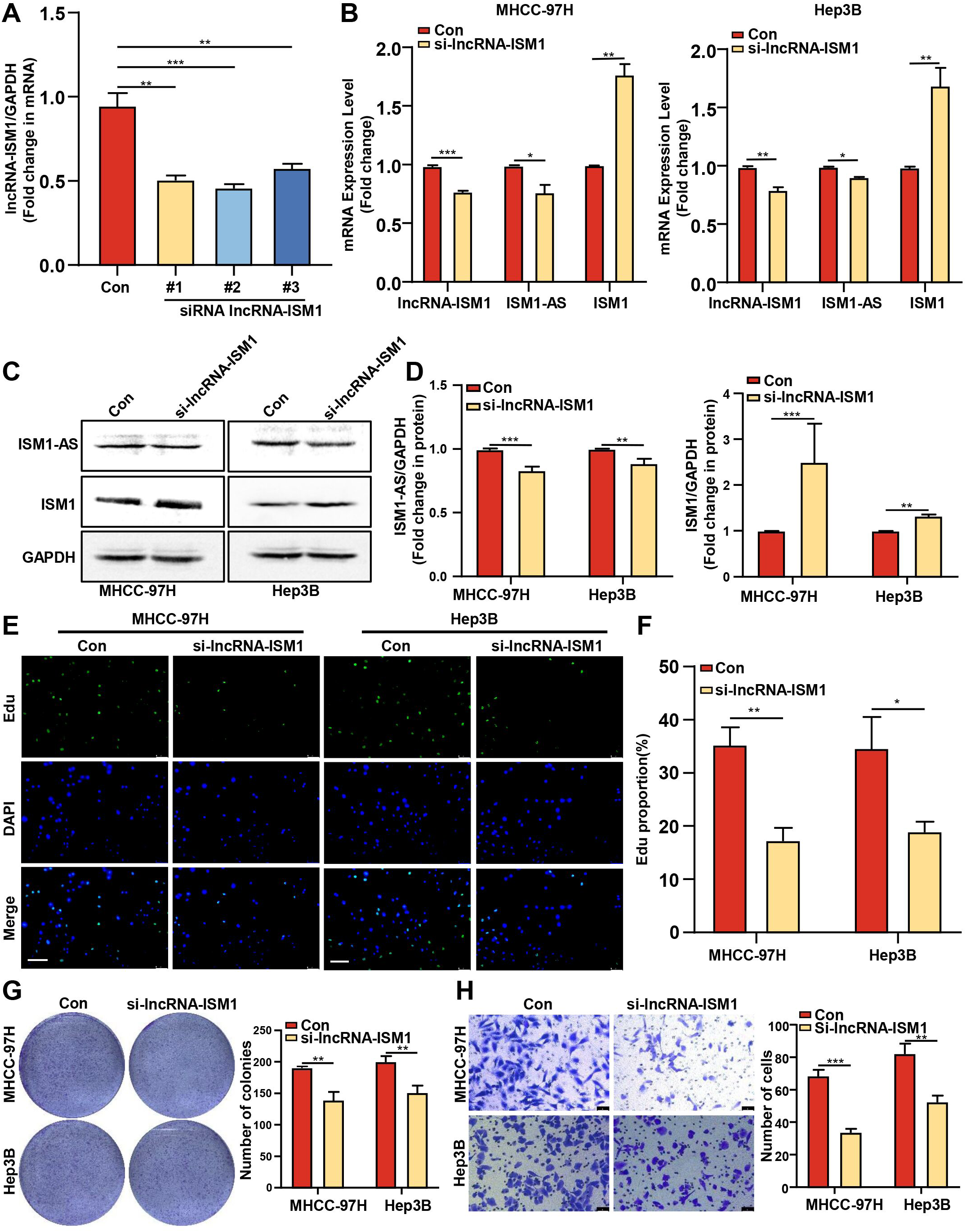
lncRNA-ISM1 regulates the proliferation and invasion of HCC cells through the ISM1-AS/ISM1 axis. MHCC-97H cells were transfected with negative control (NC) and lncRNA-ISM1 siRNA (for silencing lncRNA-ISM1) for 24 h. **(A)** qPCR was used to assess the mRNA level of lncRNA-ISM1 in the cells. The cells were subjected to **(B)** qPCR analysis of the mRNA levels of lncRNA-ISM1, ISM1-AS, and ISM1 and **(C-D)** Western blot analysis of the protein levels of lncRNA-ISM1, ISM1-AS, and ISM1. **(E–F)** EdU proliferation assays were performed using MHCC-97H and Hep3B cells. **(G)** Colony formation assays were conducted using MHCC-97H and Hep3B cells. **(H)** Transwell invasion assays were carried out using MHCC-97H and Hep3B cells. The data are presented as the mean ± standard error of the mean (mean ± SEM). n = 3. * *p* < 0.05, ** *p* < 0.01, and *** *p* < 0.001. Scale bar = 100 μm.

### 3.5 ISM1 Suppresses Glycolysis in HCC Cells via the Akt-S6 Signalling Pathway

Glycolysis plays important roles in the occurrence, development and metastasis of cancer. To further clarify the mechanism underlying the tumour-suppressive effect of ISM1, we investigated whether ISM1 expression is linked to decreased glycolysis levels. The levels of glycolysis-related proteins were assessed in MHCC-97H cells overexpressing ISM1. Compared with those in control cells, the protein levels of GLUT4, HK2, LDH and PKM2 in ISM1-overexpressing HCC cells were significantly lower **(Fig. 5A–B)**. Analyses of glucose consumption **(Fig. 5C)**, lactate production **(Fig. 5D)** and ATP content **(Fig. 5E)** also revealed that the overexpression of ISM1 reduced glucose consumption, lactate production and ATP content in HCC cells. Moreover, the overexpression of ISM1 significantly decreased the phosphorylation levels of AKT and S6, while the total levels of AKT and S6 were not affected **(Fig. 5A–B)**, suggesting that ISM1 may inhibit glycolysis in liver cancer cells by suppressing the AKT-S6 signalling pathway. To test this hypothesis, we screened siRNAs for ISM1 by qPCR **(Fig. 5F)** and used the AKT inhibitor MK2206 to inhibit the expression of AKT and S6 in MHCC-97H cells in which ISM1 was knocked down **(Fig. 5G-H)**. The results revealed that blocking AKT and S6 expression significantly reduced the levels of glycolysis-related proteins such as GLUT4, HK2, LDH and PKM2 (**Fig.** 5G–I) and simultaneously reduced glucose consumption, lactate production and ATP content. These results were verified by measuring the glucose content **(Fig. 5J)**, lactate production content **(Fig. 5K)** and ATP content analyses **(Fig. 5L)**. In conclusion, ISM1 inhibits glycolysis in HCC by suppressing AKT-S6 signalling, thereby exerting a tumour-suppressive effect.

**Fig. 5.**
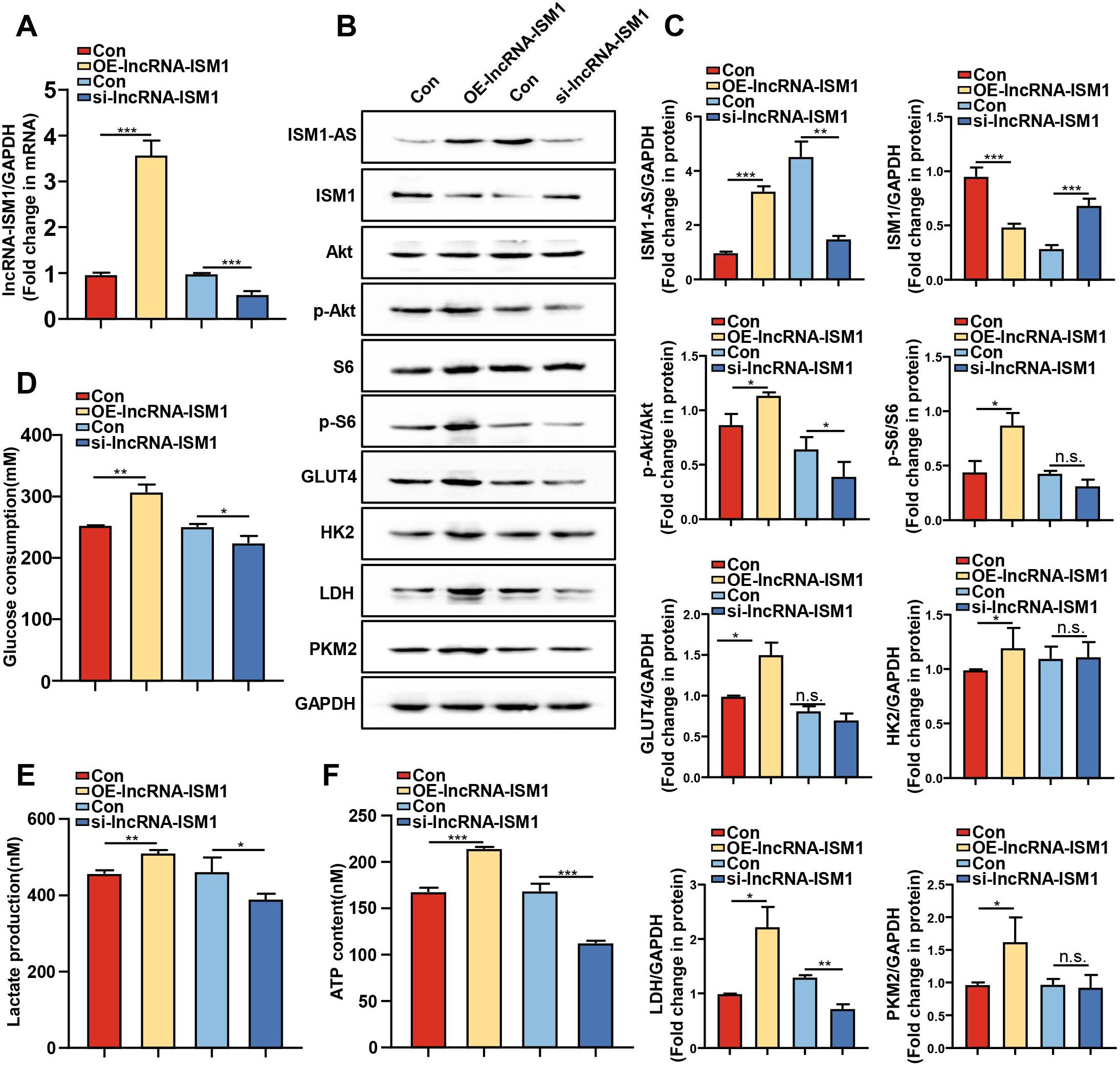
ISM1 regulates glycolysis in HCC cells through the Akt-S6 signalling pathway. An ISM1-overexpressing lentivirus was transfected into MHCC-97H cells for 24 h. **(A)** Western blotting was used to assess the protein expression of ISM1, GLUT4, HK2, LDH, and PKM2, as well as the phosphorylation of Akt and S6. The statistical results are shown in **(B)**. Glucose consumption **(C)**, lactate production **(D)**, and ATP content **(E)** were measured in MHCC-97H cells. MHCC-97H cells were transfected with negative control (NC) siRNA and ISM1 siRNA (to silence ISM1) for 24 h. **(F)** qPCR was performed to assess the mRNA level of ISM1 in the cells. MHCC-97H cells were transfected with ISM1 siRNA for 24 h and then treated with MK2206 for another 24 h. **(G)** Western blotting was used to assess the protein expression of ISM1, GLUT4, HK2, LDH, and PKM2, as well as the phosphorylation of Akt and S6. The statistical results are shown in **(H-I)**. Glucose consumption **(J)**, lactate production **(K)**, and ATP content **(L)** were measured in the cells. The data are presented as the mean ± standard error of the mean (mean ± SEM). n = 3. * *p* < 0.05, ** *p* < 0.01, and *** *p* < 0.001; n.s., not significant.

### 3.6 lncRNA-ISM1 regulates the Akt-S6 signalling pathway and glycolysis

Previous studies have shown that lncRNA-ISM1 promotes cancer through the ISM1-AS/ISM1 axis and that ISM1 inhibits glycolysis in HCC through the Akt-S6 signalling pathway. However, whether lncRNA-ISM1 also regulates HCC glycolysis through the Akt-S6 signalling pathway remains unclear. Therefore, we transfected lncRNA-ISM1 overexpression and knockdown vectors into MHCC-97H cells **(Fig. 6A)**. The results revealed that the overexpression of lncRNA-ISM1 led to an increase in the level of ISM1-AS and a decrease in the level of ISM1, which was consistent with previous results. In addition, the overexpression of lncRNA-ISM1 significantly increased the phosphorylation levels of Akt and S6 in the cells, while the total levels of Akt and S6 were not affected (**Fig.** 6B–C). Moreover, the protein levels of GLUT4, HK2, LDH and PKM2 in HCC cells also increased significantly **(Fig. 6B–C)**. Consistent with these findings, the results of glucose content **(Fig. 6D)**, lactate production **(Fig. 6E)** and ATP content **(Fig. 6F)** assays revealed that after the overexpression of lncRNA-ISM1, glucose consumption, lactate production and ATP content increased, whereas lncRNA-ISM1 knockdown resulted in the opposite trend. These results confirm that lncRNA-ISM1 regulates glycolysis in HCC cells through the Akt-S6 signalling pathway.

**Fig. 6.**
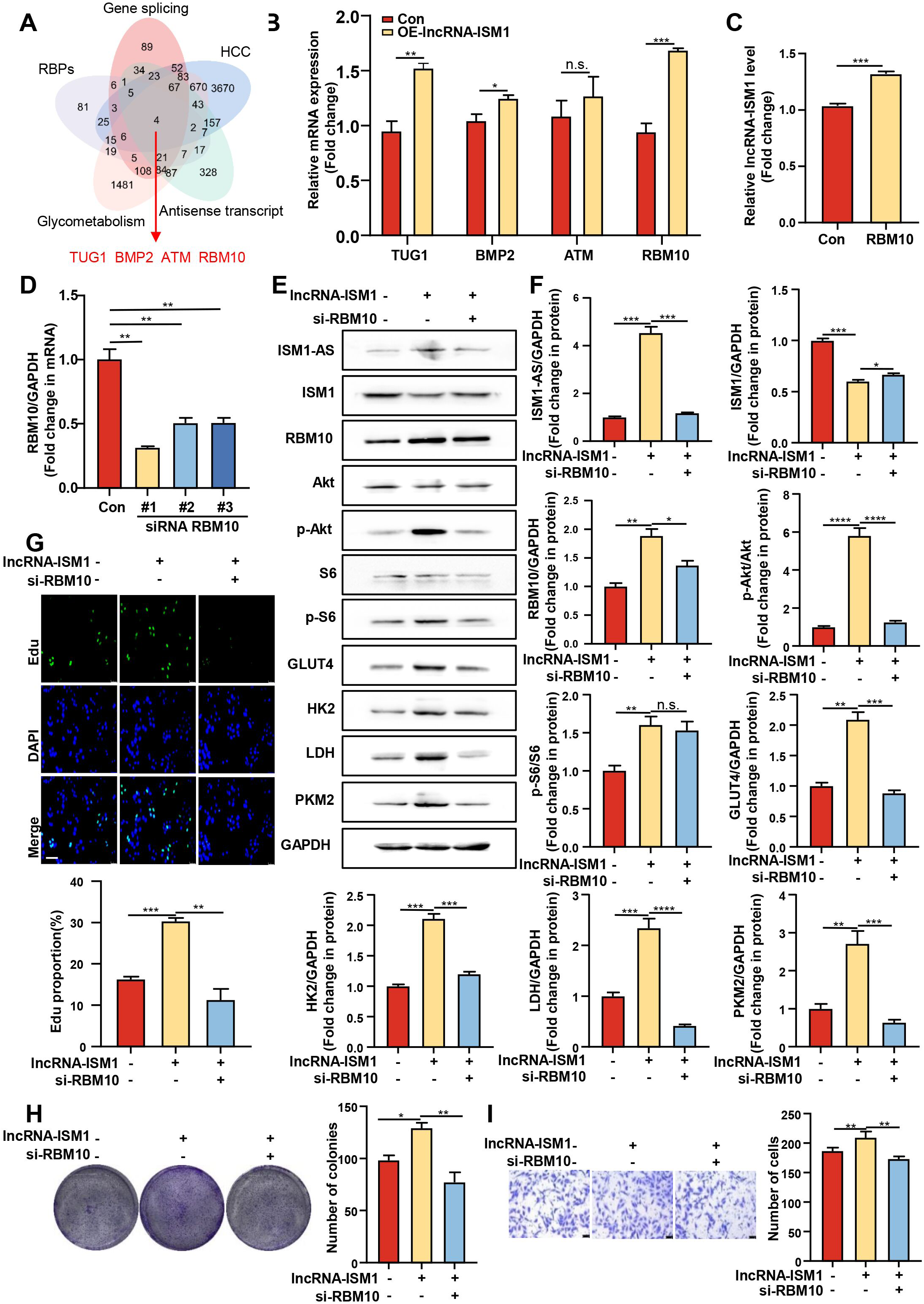
lncRNA-ISM1 promotes glycolysis in HCC cells through Akt-S6 signalling pathway activation by altering the ISM1-AS/ISM1 balance. MHCC-97H cells were transfected with either a lncRNA-ISM1-overexpressing lentivirus or lncRNA-ISM1 siRNA. **(A)** qPCR was used to assess the mRNA level of lncRNA-ISM1 in the cells. **(B)** Western blotting was performed to assess the protein expression of ISM1-AS, ISM1, GLUT4, HK2, LDH, and PKM2, as well as the phosphorylation of Akt and S6. The statistical results are shown in **(C)**. Glucose consumption **(D)**, lactate production **(E)**, and ATP content **(F)** were measured in MHCC-97H cells. The data are presented as the mean ± standard error of the mean (mean ± SEM). n = 3. * *p* < 0.05, ** *p* < 0.01, and *** *p* < 0.001; n.s., not significant.

### 3.7 lncRNA-ISM1 Interacts with RBM10 to Suppress the ISM1-AS/ISM1 Axis and Alleviates HCC Cell Proliferation and Migration

The current data preliminarily reveal the mechanism by which lncRNA-ISM1 promotes the progression of HCC through AKT-S6 signalling pathway activation through the ISM1-AS/ISM1 axis. Therefore, the target genes of lncRNA-ISM1 may serve as promising targets for HCC treatment. We used the GeneCards database (https://www.genedcards.org) to screen the target genes of lncRNA-ISM1 online with the keywords “HCC”, “Gene splicing”, “RBPs”, “Antisense transcript”, and “glycometabolism”. The genes predicted by the database were intersected using a Venn diagram, and the following 4 genes were screened out: TUG1, BMP2, ATM, and RBM10 **(Fig. 7A)**. After lncRNA-ISM1 was overexpressed in MHCC-97H cells, the expression of RBM10 was the most significantly upregulated **(Fig. 7B)**. RIP experiments confirmed that RBM10 binds to lncRNA-ISM1 **(Fig. 7C)**. On this basis, we speculate that RBM10 may interact with lncRNA-ISM1 and regulate its downstream molecules. Next, we screened siRNAs for RBM10 by qPCR **(Fig. 7D)** and knocked down the expression of RBM10 in MHCC-97H cells overexpressing lncRNA-ISM1. The results showed that knocking down RBM10 reversed the effects of lncRNA-ISM1 on HCC cell glycolysis and progression. Western blot analysis revealed that knocking down RBM10 in MHCC-97H cells inhibited the protein expression of ISM1-AS and promoted the protein expression of ISM1; moreover, it significantly reduced the protein levels of phosphorylated AKT, S6, GLUT4, HK2, LDH, and PKM2 **(Fig. 7E–F)**. In addition, decreases in cell proliferation **(Fig. 7G-H)** and invasion **(Fig. 7I)** were observed in MHCC-97H cells after RBM10 knockdown. These results indicate that knocking down RBM10 expression can inhibit the proliferation and migration of HCC cells by blocking the regulation of the ISM1-AS/ISM1 axis by lncRNA-ISM1.

**Fig. 7.**
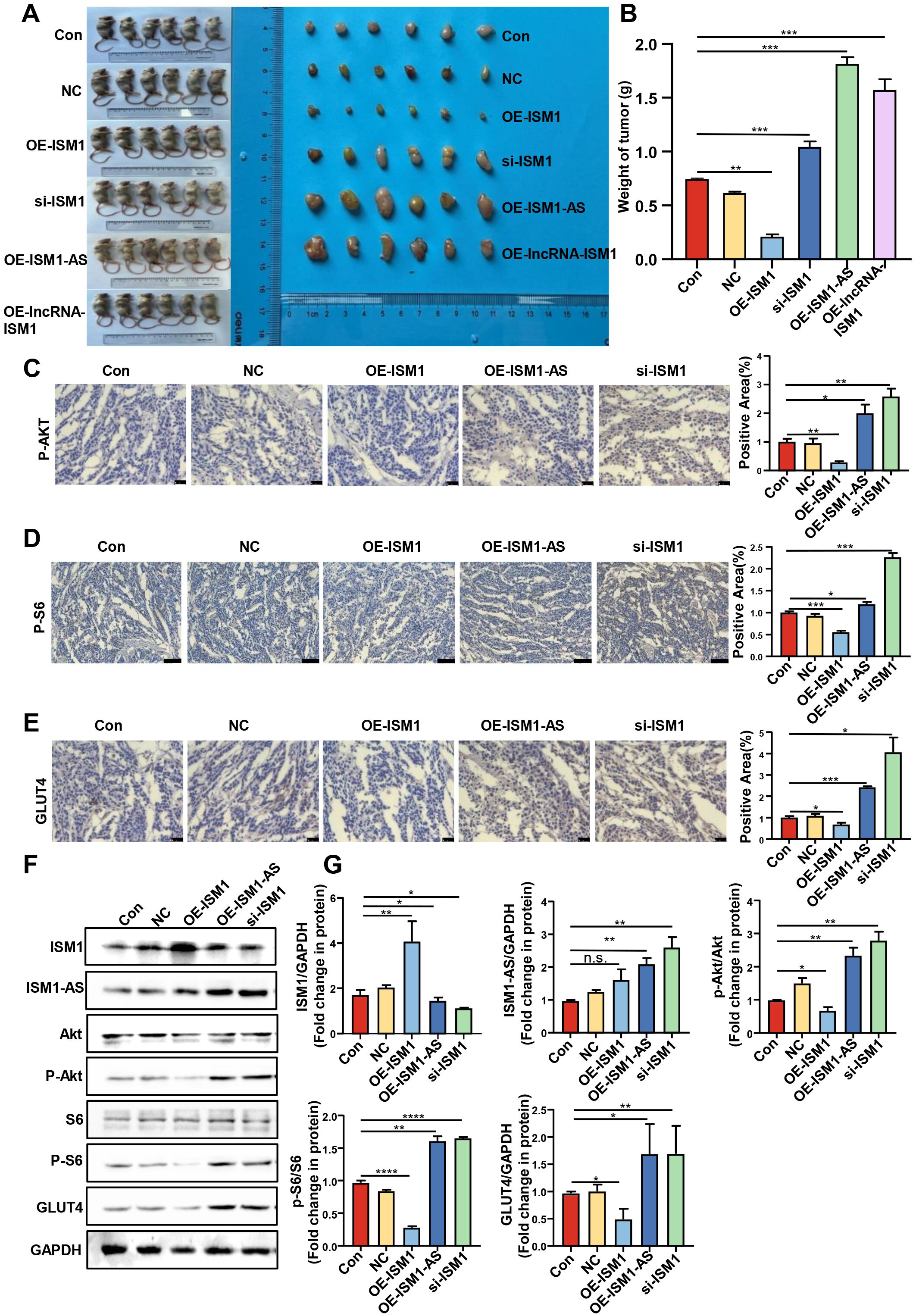
lncRNA-ISM1 and RBM10 interact to regulate glycolysis and cell progression in HCC cells. **(A)** Datasets related to HCC, gene splicing, RBPs, glycometabolism, and antisense transcripts were downloaded from the GeneCards database (https://www.genedcards.org), and a Venn diagram was constructed to show the intersections. **(B)** qPCR was used to assess the mRNA levels of TUG1, BMP2, ATM, and RBM10 in cells overexpressing lncRNA-ISM1. **(C)** RNA immunoprecipitation assays were performed using an anti-RBM10 antibody in MHCC-97H cells. **(D)** After MHCC-97H cells were transfected with RBM10 siRNA for 24 h, qPCR was used to assess the mRNA level of RBM10 in the cells. **(E)** MHCC-97H cells overexpressing lncRNA-ISM1 were transfected with RBM10 siRNA, and Western blotting was used to assess the protein expression levels of ISM1-AS, ISM1, RBM10, GLUT4, HK2, LDH, and PKM2, as well as the phosphorylation levels of Akt and S6. The statistical results are shown in **(F)**. **(G)** EdU-based proliferation assays were performed using MHCC-97H cells treated as described in **(E)**. **(H)** Colony formation assays were performed using MHCC-97H cells treated as described in **(E)** for 14 days. **(I)** Transwell invasion assays were performed using MHCC-97H cells treated as described in **(E)** for 24 h. The data are presented as the mean ± standard error of the mean (mean ± SEM). n = 3. * *p* < 0.05, ** *p* < 0.01, *** *p* < 0.001, and **** *p* < 0.0001. Scale bar = 100 μm.

### 3.8 In Vivo Study of Tumour Growth Regulation by lncRNA-ISM1 and ISM1

To verify the effects of lncRNA-ISM1 and ISM1 on HCC in vivo, we subcutaneously injected MHCC-97H cells with stable ISM1 overexpression, ISM1 knockdown, lncRNA-ISM1 overexpression, or ISM1-AS overexpression or control cells into nude mice. The expression levels of ISM1 and ISM1-AS proteins in tissues were assessed using immunofluorescence experiments **(Fig. S4A–D)**. Four weeks after injection, the volume and weight of HCC tumours were evaluated. Compared with those in the control group, the volume and weight of tumours in nude mice injected with ISM1-overexpressing cells were significantly reduced, indicating that the overexpression of ISM1 inhibited the growth of HCC tumours in vivo **(Fig. 8A–B)**. In contrast, the volume and weight of tumours in nude mice injected with ISM1 knockdown, ISM1-AS overexpression, or ISM1-overexpressing cells significantly increased **(Fig. 8A-B)**. Immunohistochemical analysis of the tumour tissues of transplanted mice revealed that the overexpression of ISM1 significantly reduced the protein levels of P-AKT, P-S6, and GLUT4, whereas the overexpression of ISM1-AS and knockdown of ISM1 increased the protein levels of P-AKT, P-S6, and GLUT4 (**Fig.** 8C–E). Western blot analysis also revealed that the overexpression of ISM1 significantly decreased the phosphorylation levels of AKT and S6 and the protein level of GLUT4, whereas the overexpression of ISM1-AS and knockdown of ISM1 increased the phosphorylation levels of AKT and S6 and the protein level of GLUT4 **(Fig. 8F–G)**, findings that are consistent with the results of the in vitro experiments. Our results indicate that regulating the expression of lncRNA-ISM1 and ISM1 can affect the growth of HCC in vivo.

**Fig. 8.**
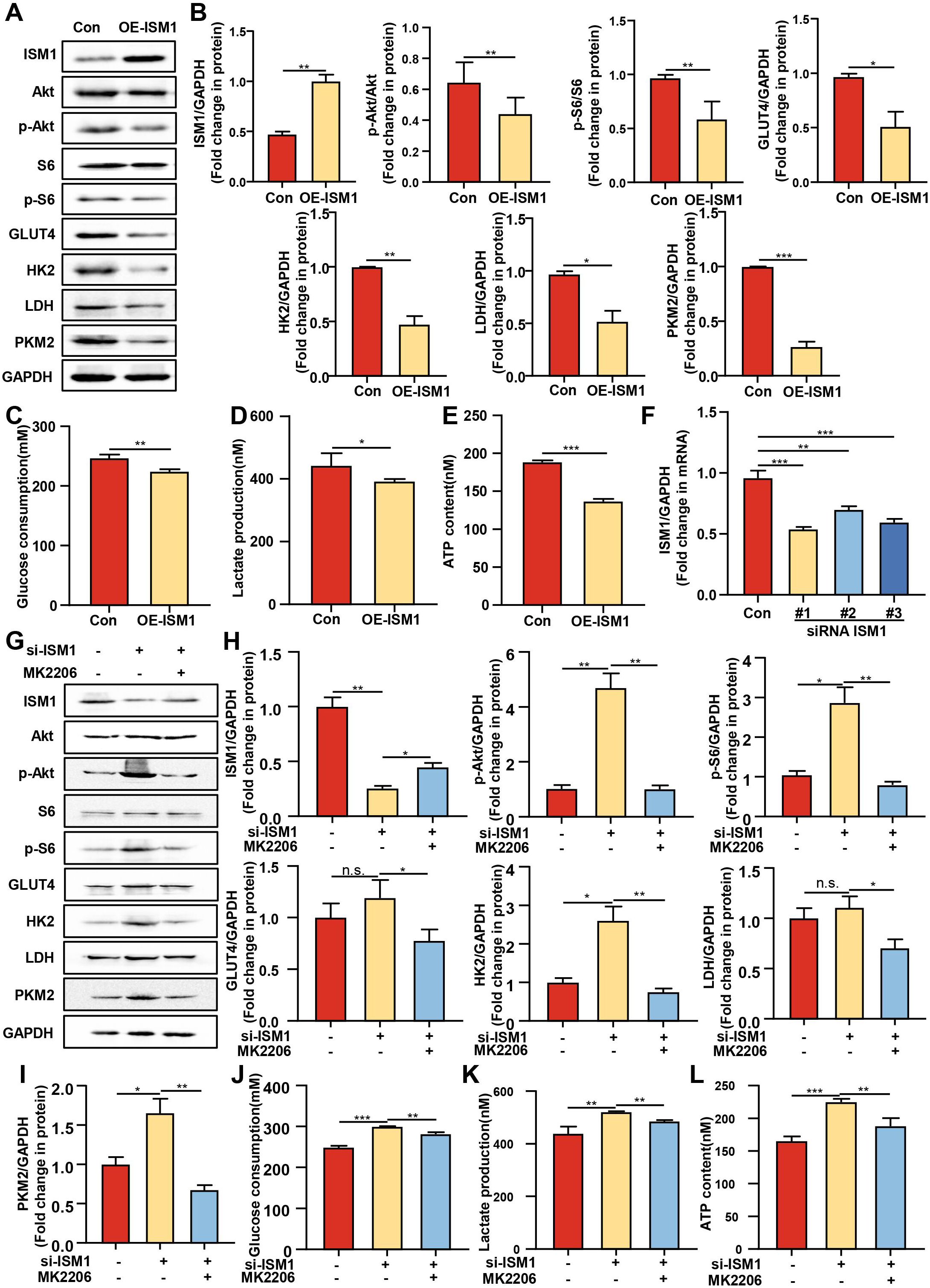
ISM1 promotes HCC growth in mice via Akt-S6 signalling pathway activation. BALB/c nude mice (6 weeks old, weighing 18–22 g) were subcutaneously inoculated with human HCC MHCC-97H cells (2×10⁶ cells/mouse) on the right posterior back. The mice were divided into a control group, an NC group, an ISM1-overexpression group, an ISM1-knockdown group, a lncRNA-ISM1-overexpression group, and an ISM1-AS-overexpression group. The mice were observed for 4 weeks after inoculation, after which the tumours were dissected, photographed, and weighed. Representative photographs of tumour appearance and size on the backs of nude mice are shown in (A). **(B)** An analysis of tumour size is presented. Immunohistochemistry was used to assess the expression levels of P-AKT **(C)**, P-S6 **(D)**, and GLUT4 in mouse HCC tissues. **(F)** Western blotting was performed to assess the protein expression levels of ISM1, ISM1-AS, and GLUT4, as well as the phosphorylation levels of Akt and S6. The statistical results are shown in **(G)**. The data are presented as the mean ± standard error of the mean (mean ± SEM). n = 3-6. * *p* < 0.05, ** *p* < 0.01, *** *p* < 0.001, and **** *p* < 0.0001. Scale bar = 25 μm.

## 4. Discussion

Hepatocellular carcinoma (HCC) is among the most common malignant tumours in clinical practice. Although significant progress has been made in the clinical treatment of HCC over the past few decades, the exact mechanisms underlying its initiation and development remain incompletely understood. In this study, we revealed that the interaction between lncRNA-ISM1 and RBM10 regulates the ISM1-AS/ISM1 axis and suppresses the AKT-S6 signalling pathway, thereby inhibiting the growth and invasion of hepatocellular carcinoma.

ISM1 was initially recognized for its role as an angiogenesis inhibitor in various tumours [36]. In recent years, owing to its mutation or aberrant expression in multiple cancers, the function of ISM1 in tumorigenesis has garnered increasing attention [37]. For example, compared with that in normal colonic mucosal epithelial tissues, ISM1 mRNA expression is upregulated in colon adenocarcinoma cell lines, whereas the targeting of ISM1 by miR-1307-3p inhibits ISM1 expression, thereby suppressing the proliferation of colon adenocarcinoma cells [19]. Wu et al. demonstrated that ISM1 can promote the migration and proliferation of colorectal cancer cells[18]. Moreover, ISM1 can serve as a tumour biomarker, as it is highly expressed in aldosterone-producing adenomas with CTNNB1 mutations [38]. ISM1 methylation can also serve as a biomarker for the survival of women with invasive lobular breast cancer [39]. Similarly, ISM1 is differentially expressed in uveal melanoma [40] and cholangiocarcinoma [41]. However, the role of ISM1 in the pathogenesis of hepatocellular carcinoma (HCC) remains unclear. In this study, we revealed that ISM1 has a tumour-suppressive function in HCC development. Data mining from the TCGA database revealed that ISM1 expression is lower in HCC tumour tissues than in normal tissues. Importantly, experimental studies confirmed that compared with its expression in adjacent nontumour tissues and cells, ISM1 expression is significantly downregulated in clinical HCC tumour tissues and HCC cell lines. In HCC cells, we also demonstrated that the overexpression of ISM1 inhibited cell proliferation and invasion, whereas the knockdown of ISM1 significantly promoted proliferation and invasion. These results were validated in tumour xenograft experiments in vivo. For the first time, we revealed the tumour-suppressive function of ISM1 in regulating HCC progression.

In this study, we propose a novel potential mechanism by which ISM1 inhibits the proliferation and invasion of HCC through the suppression of the AKT-S6 signalling pathway. The PI3K/Akt/mTOR signalling pathway has emerged as a central hub for multiple oncogenic signalling pathways and is an attractive drug target in cancer therapy. It can trigger tumour cell motility and invasion. Both mTOR complexes promote tumour cell migration in different ways: mTORC1 induces tumour cell migration and invasion through its downstream targets, i.e., p70 ribosomal protein S6 kinase and eukaryotic initiation factor 4E-binding protein [42]. In this study, we revealed that the knockdown of ISM1 activates the AKT-S6 signalling pathway, whereas blocking AKT expression reduces the proliferation, migration, and invasion of HCC cells induced by ISM1 knockdown. Previous studies have reported that ISM1 regulates signalling during zebrafish embryonic development through the Wnt/β-catenin and Nodal pathways[42, 43]. Nevertheless, whether ISM1 inhibits HCC progression by regulating other signalling pathways remains to be further investigated.

Recent studies have shown that alternative splicing plays a crucial role in cancer and interacts with other molecular mechanisms to jointly promote cancer initiation and progression [23]. The role of alternative splicing in cancer has attracted increasing attention because it can generate multiple protein variants that play important roles in tumour initiation, development, and drug resistance. In cancer, dysregulated alternative splicing is considered a key factor driving tumour progression. Therefore, targeting mis-spliced genes (i.e., genes with significantly altered splicing in cancer) has become a powerful strategy for cancer therapy [44, 45]. In April 2022, GenBank released a new splice variant of ISM1 that encodes a protein lacking the AMOP domain. This new mutation may affect the ability of ISM1 to regulate cellular behaviour. Interestingly, our study revealed that ISM1 ameliorates the carcinogenic-promoting effect of ISM1-AS on liver cancer and that this process plays a key role in regulating the proliferation and invasion of HCC. However, the role of ISM1-AS in HCC remains unclear, and the specific mechanisms through which it induces ISM1 accumulation in HCC cells require further elucidation.

In cancer research, the aberrant expression of antisense transcripts is closely associated with tumour initiation, progression, and prognosis, including in hepatocellular carcinoma [46]. Recent studies have shown that antisense transcripts play a role in cancer development by suppressing the splicing and translation of their parental genes, suggesting that increasing the number of antisense transcripts that target oncogenes or silencing those that target tumour suppressor genes may serve as potential strategies for cancer treatment [47]. Antisense transcripts play a crucial role in gene expression regulation by binding to sense mRNA, affecting its stability, translation efficiency, or alternative splicing, thereby modulating gene expression. Specifically, during alternative splicing, antisense transcripts can bind to mRNA, influencing spliceosome assembly or the binding of splicing factors and thereby altering splicing outcomes [48]. In this study, our data demonstrate that the overexpression of lncRNA-ISM1 leads to increased ISM1-AS levels and decreased ISM1 levels, promoting HCC cell proliferation and migration, whereas the knockdown of lncRNA-ISM1 reverses the effects of lncRNA-ISM1 on ISM1-AS and ISM1 activity and inhibits HCC cell proliferation and invasion. These findings highlight the potential value of lncRNA-ISM1 as a key regulatory molecule of ISM1-AS expression and as a therapeutic target and diagnostic biomarker for HCC.

On the basis of the aforementioned findings, we identified RBM10 as a target gene of lncRNA-ISM1 through the GeneCards database and confirmed the interaction between RBM10 and lncRNA-ISM1 using RIP experiments. RBM10 is an RNA-binding protein that appears to exert its oncogenic effects by binding to lncRNA-ISM1. However, in some cancer cells, RBM10 also has strong antitumour properties [49]. Nevertheless, the mechanisms underlying the function of RBM10 remain incompletely understood. In this study, we revealed that RBM10 promotes the proliferation and invasion of HCC cells and that knocking down RBM10 in cells overexpressing lncRNA-ISM1 reverses the effects of lncRNA-ISM1 on ISM1-AS and ISM1, thereby inhibiting HCC cell proliferation and invasion. This may help explain the role of RBM10 in cancer therapy.

Our findings highlight the potential clinical significance of the pathological correlations among lncRNA-ISM1, RBM10, and ISM1-AS. The expression level of ISM1 holds promise as a potential biomarker for HCC diagnosis and prognosis. Additionally, blocking the interaction between lncRNA-ISM1 and RBM10 may provide a novel strategy for treating HCC, as inhibiting their binding is crucial for reducing ISM1-AS, promoting ISM1 accumulation, and suppressing HCC glycolysis and cellular progression. Such intervention can disrupt downstream cell signalling activity, thereby effectively inhibiting the development of liver cancer. Currently, drugs targeting lncRNA-ISM1 are typically antisense oligonucleotides (ASOs) [50]. Although there are currently no drugs specifically targeting lncRNA-ISM1, future research may lead to the development of ASO drugs targeting ISM1.

## 5. Conclusion

In summary, our current study reveals a novel function and mechanism of lncRNA-ISM1 in HCC progression, namely, by interacting with RBM10 to regulate the expression of ISM1-AS and ISM1, thereby activating the AKT-S6 signalling pathway and ultimately promoting the growth and metastasis of HCC.

## Abbreviations

HCC: Hepatocellular Carcinoma
ISM1: Isthmin-1
AS: Alternative splicing
LncRNA: Long non-coding RNA
GLUT4: Glucose transporter 4
HK2: Hexokinase 2
LDH: Lactate Dehydrogenase
PKM2: Pyruvate Kinase M2 Isoform
EDU: 5-Ethynyl-2’-deoxyuridine
RBP: RNA binding protein
ATP: Adenosine Triphosphate
TUG1: taurine up - regulated gene 1
BMP2: Bone Morphogenetic Protein 2
ATM: Ataxia-Telangiectasia Mutated
RBM10: RNA Binding Motif Protein 10
AKT: Protein Kinase B
S6: Ribosomal protein S6

## Ethics approval and consent to participate

All human samples used in experiments were approved by the Ethics Committee of Xi’an No.3 Hospital, the Affiliated Hospital of Northwest University (No. SYLL-2025-101). All experiment procedures and animal caring were in accordance with the institutional ethics directions for animals-related experimental processes.

## Author Contributions

**Man Li**, and **Deqing Huang**: Writing – original draft, Writing – review & editing, Visualization. **Yuanyuan Ren**: Writing – review & editing, Visualization. **Zhen Wang**: Validation. **Yujia Li**: Writing – review & editing. **Weichen Zuo**: Writing – review & editing. **Yijie Li**: Writing – review & editing. **Yanyan Jin**: Writing – review & editing. **Yuyan Xiong**: Writing – revision and finalization, Supervision.

